# Mapping lignification dynamics with a combination of chemistry, data segmentation and ratiometric analysis

**DOI:** 10.1101/2021.06.21.449296

**Authors:** Oriane Morel, Cedric Lion, Godfrey Neutelings, Jonathan Stefanov, Fabien Baldacci-Cresp, Clemence Simon, Notburga Gierlinger, Christophe Biot, Simon Hawkins, Corentin Spriet

## Abstract

This article describes a new methodology for detailed mapping of the lignification capacity of plant cell walls that we have called “REPRISAL” for **REP**orter **R**atiometrics **I**ntegrating **S**egmentation for **A**nalyzing **L**ignification. REPRISAL consists of the combination of three separate approaches. In the first approach, H*, G* and S* monolignol chemical reporters, corresponding to *p*-coumaryl alcohol, coniferyl alcohol and sinapyl alcohol, are used to label the growing lignin polymer in a fluorescent triple labelling strategy based on the sequential use of 3 main bioorthogonal chemical reactions. In the second step, an automatic parametric and/or artificial intelligence (AI) segmentation algorithm is developed that assigns fluorescent image pixels to 3 distinct cell wall zones corresponding to cell corners (CC), compound middle lamella (CML) and secondary cell walls (SCW). The last step corresponds to the exploitation of a ratiometric approach enabling statistical analyses of differences in monolignol reporter *distribution* (ratiometric method 1) and *proportions* (ratiometric method 2) within the different cell wall zones. In order to demonstrate the potential of REPRISAL for investigating lignin formation we firstly describe its use to map developmentally-related changes in the lignification capacity of WT Arabidopsis interfascicular fiber cells. We then show how it can be used to reveal subtle phenotypical differences in lignification by analyzing the Arabidopsis *prx64* peroxidase mutant and provide further evidence for the implication of the AtPRX64 protein in floral stem lignification. Finally, we demonstrate the general applicability of REPRISAL by using it to map lignification capacity in poplar, flax and maize.

## Introduction

Lignin is a phenolic polymer found in the walls of certain plant cells where it makes up approximately 25-30 % of the dry weight. Together with cellulose and xylan, it is one of the major constituents of wood and is the second most abundant plant polymer after cellulose on Earth. Both the quantity and the structure of lignin present in plant cell walls have a major effect on the mechanical and chemical properties of numerous plant-derived resources/products including wood, paper and textiles (Huis et al., 2012). Lignin is also a major factor responsible for the recalcitrance of ligno-cellulose biomass in the industrial production of bio-ethanol for energy. For plants themselves, the presence of lignin in the cell wall contributes positively to mechanical support, facilitates water transport in xylem via its hydrophobicity and helps to protect them against pathogens and physical injury (Boerjan et al., 2003). The appearance of lignin during evolution has played a major role in ensuring the survival of plants on dry land. The lignin polymer is also an important carbon sink and the formation of this polymer in the wood cells of long-lived tree species therefore contributes positively in the struggle against climate change.

Chemically, lignin is formed by the non-enzymatic polymerisation of phenoxy radicals mainly derived from three main monomers: *para*-coumaryl alcohol, coniferyl alcohol and sinapyl alcohol that are collectively referred to as ‘monolignols’ (Ralph et al., 2004). Once incorporated into the polymer, these molecules form the hydroxyphenyl (H), guaiacyl (G) and sinapyl (S) monomeric units. Lignin can also contain C (catechyl)-units derived from caffeyl alcohol in orchids and the cactaceae families (Chen et al., 2012) as well as tricin flavonoids and ferulates in grasses (Grabber et al., 2000; del Río et al., 2012) or hydroxystilbenes in palm fruit (del Río et al., 2017). The amount of lignin and the relative proportion of the different monomers in the cell wall varies according to the botanical group (e.g. gymnosperm lignin contains no/very few S units compared to angiosperm lignin which contains far higher quantities of S units); the organ (leaves are far less lignified than stems/roots), the tissue/cell type (e.g. in Arabidopsis, xylem vessels have a lower S/G ratio than fibers), and the cell wall layer (cells with only primary cell walls are generally non-lignified whereas cells with secondary cell walls become lignified) (Campbell and Sederoff, 1996). Monolignols are enzymatically synthesized via the phenylpropanoid pathway and are then exported into the developing cell wall across the plasma membrane. Several hypotheses including the involvement of monolignol specific transporters, vesicle-related transport and passive diffusion have all been advanced to explain monolignol transport, but the actual mechanism(s) involved have not as yet been categorically determined and further research is necessary on this point (Perkins et al., 2019). Once in the cell wall compartment, monolignols are oxidized by cell wall located laccases and/or peroxidases before undergoing polymerization into the growing lignin polymer (Wang et al., 2013).

In order to understand the dynamics of lignification at the plant or organ scale, different analytical techniques have been developed over the last decades (Lupoi et al., 2015). They are generally adapted to the species studied (woody or not) and to the quantities of lignins present in their tissues. For example, the method using acetyl bromide solubilization of lignins (Johnson et al., 1961) is well suited for herbaceous species while the gravimetric Klason method (Effland, 1977) is more suited for woody species. There are also different methods for determining the H, G, and S subunit composition and the S/G ratio of lignins. Historically, thioacidolysis (Lapierre et al., 1986) and nitrobenzene oxidation (Billa and Monties, 1995) methods were used, and are now gradually replaced by pyrolysis coupled with GC/MS (Wagner et al., 2007) or NMR (Mansfield et al., 2012).

All these approaches provide relevant information on the different tissues/organs studied but are destructive, meaning that it is not possible to get information regarding quantities and compositions at the cellular level. Therefore, histochemical approaches are usually associated with these global analyses such as phloroglucinol staining (Wiesner reaction), which specifically reveals the cinnamaldehyde functions of S and G unit derivatives by producing a red chromogen whose intensity depends on the quantities (Clifford, 1974). The Maüle test uses potassium permanganate and hydrochloric acid to transform guaiacyl and syringyl residues into catechols, which are transformed into orange-brown quinones for G lignins and purple-red quinones for G-S lignins (Day et al., 2005). Other approaches exploit the autofluorescence of lignins (Donaldson, 2001) or the possibility to visualize them using fluorescent compounds such as auramine (Pesquet et al., 2005) and acriflavine (Donaldson, 2001). All these techniques therefore allow to obtain relatively fine spatial information, but not a clear view on the dynamics of lignification since visualization of the active zones of monolignol polymerization in tissues is lacking.

To overcome this intrinsic drawback, the bioorthogonal chemical reporter strategy has gained increased attention in bioimaging because it enables the study of *de novo* lignin formation in living organisms. In this two-step strategy, a synthetic derivative (the chemical reporter) of a monolignol is first introduced into the cell wall where it becomes oxidized by cell wall located peroxidases and/or laccases and incorporated into the growing lignin polymer. The reporter carries a specific chemical group that enables subsequent interaction with a corresponding probe *via* a specific bioorthogonal chemical reaction. To enable integration, the group must be small and innocuous so as to minimize its impact on the physicochemical properties of the reporter and allow its recognition by the enzymatic machinery (Rigolot et al., 2021). Once metabolic incorporation has occurred, the reporter group must then react specifically and efficiently with an exogenous molecular probe (e.g., an adequately functionalized flurophore) while being chemically inert to the surrounding biological environment. Different bioorthogonal reactions have been developed over the past 20 years and mainly applied to glycan and protein labelling (Wallace and Chin, 2014; Palaniappan and Bertozzi, 2016). In 2014, Ralph and Zhu’s teams simultaneously published the first use of coniferyl alcohol analogues as a lignin chemical reporter in a mono-labelling strategy (Bukowski et al., 2014; Tobimatsu et al., 2014). More recently our group elaborated a double-, and then triple-labelling strategy to visualize active lignification areas in plants. For this we designed three chemical reporters (G*, H* and S*) mimicking the three main monolignols (G, H and S), tagged with an alkyne group, an azide function and a methylcyclopropene moiety, respectively) and exploited the three main bioorthogonal reactions, namely the copper-catalyzed alkyne-azide cycloaddition (CuAAC), the strain-promoted azide-alkyne cycloaddition (SPAAC) and the inverse electronic demand Diels-Alder cycloaddition (IEDDA) (Lion et al., 2017; Simon et al., 2018). The triple labelling strategy was then used to characterize the dynamics of lignification in flax (*Linum usitatissimum* L.) and we also showed that the method could be transposed to other plant species (*Arabidopsis thali*ana, tobacco and poplar). This technology provided a powerful new approach to investigate lignification at the cell wall/cell wall layer scale in different plant species. However, while a simple visual inspection of different samples is sufficient to detect “differences” in monolignol reporter incorporation profiles, often the sheer complexity of the data obtained makes it more or less impossible to qualify and/or quantify all but the most obvious differences (e.g., lignified *vs* non-/poorly-lignified walls) thereby limiting the biological potential of the chemical reporter approach.

We therefore decided to develop a segmentation approach that when combined with the chemical reporter strategy can be used for statistical determination of relative reporter incorporation into different cell wall zones based upon a ratiometric method. In this paper we present this combined methodology that we have called “REPRISAL” for **REP**orter **R**atiometrics **I**ntegrating **S**egmentation for **A**nalyzing **L**ignification. We describe the development of REPRISAL and demonstrate how it can be used for detailed mapping of lignification in WT Arabidopsis, in the *prx64* peroxidase mutant as well as in poplar, flax and maize.

## Results and discussion

REPRISAL consists of three separate approaches: 1) bioorthogonal labeling of lignin with the three chemical reporters, 2) parametric or artificial intelligence (AI) segmentation of labelled cell walls and 3) ratiometric analysis of fluorescence signal intensity (Figure 1). Each approach is described in more detail below.

**Figure 1.**
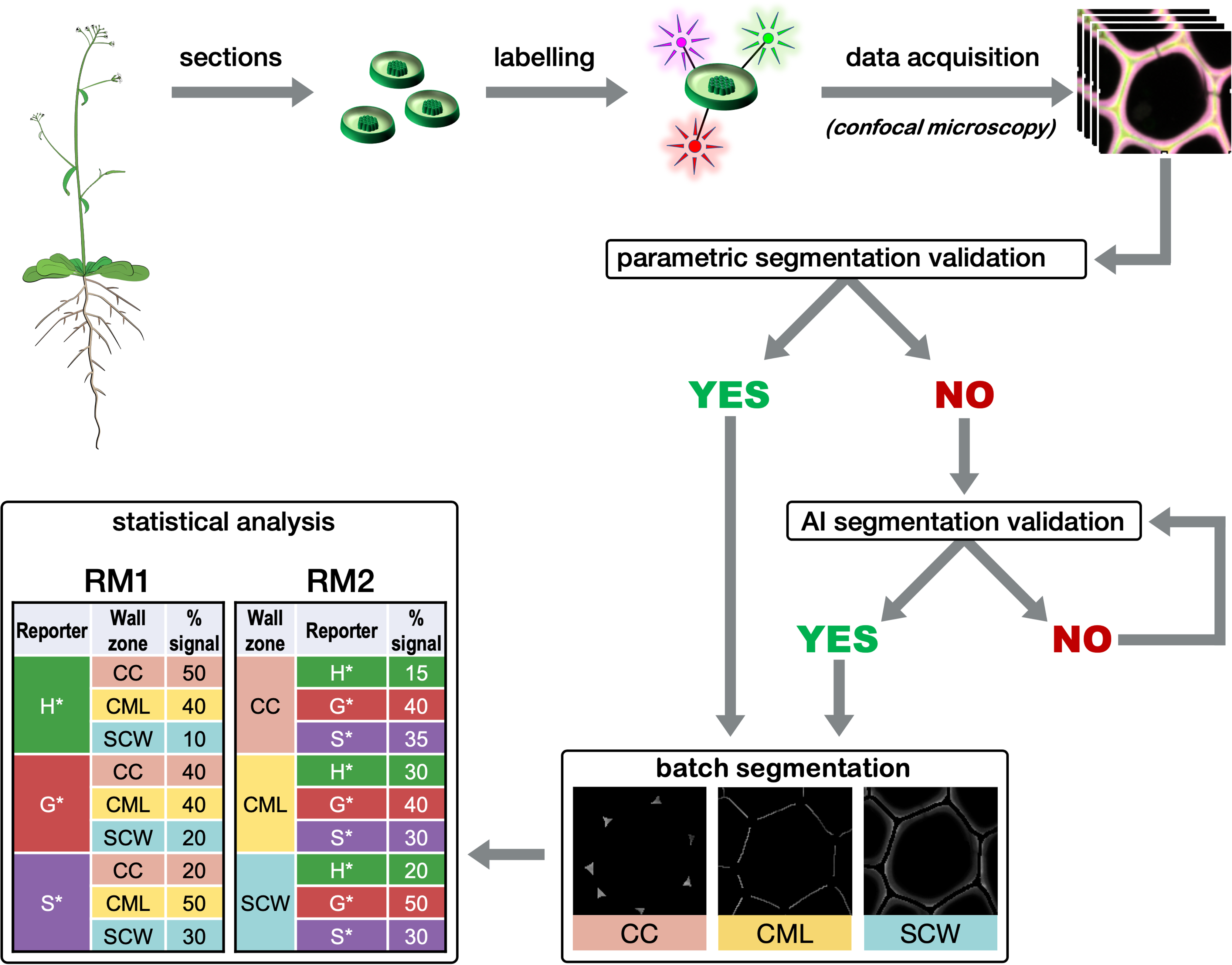
Global procedure workflow. Plant stem transversal cross-sections are incubated with H*, G* and S* monolignol reporters that become metabolically incorporated into the growing lignin polymer. Incorporated reporters are identified via a bioorthogonal chemistry strategy specifically linking each type of reporter to a different fluorescent probe allowing imaging by confocal microscopy (see Fig. 2). Following image acquisition, a first representative image of the series is selected and the most suitable segmentation method (parametric or AI) is chosen and applied automatically to all acquired images. The data is then extracted for 3 different cell wall zones (CC: cell corner, CML: compound middle lamella, SCW: secondary cell wall) and relative reporter distribution and proportion determined by ratiometric method 1 (RM1) and ratiometric method 2 (RM2). Please see text for more details

### 1. Bioorthogonal labeling and imaging

Three synthetic monolignol surrogates H*, G* and S* (the chemical reporters) equipped with click-reactive tags (Figure 2) were metabolically incorporated into *A. thaliana* stem cross-sections *ex vivo* using our previously reported triple bioorthogonal labelling methodology (Simon et al., 2018). Each incorporated reporter was then specifically linked to a fluorescent probe via a specific bioorthogonal ligation reaction thereby enabling subsequent fluorescence visualization and identification of the tagged substrates. H*-units were labelled with DBCO-Rhodamine Green (SPAAC bioorthogonal reaction), G*-units were labelled with Azide Fluor 545 (CuAAC bioorthogonal reaction), and S*-units were labelled with Tetrazine-Cy5 (IEDDA bioorthogonal reaction). Samples were then visualized by confocal microscopy to produce a set of images based on 3-channel detection (green for H*, red for G* and magenta for S*) (Figure 2).

**Figure 2.**
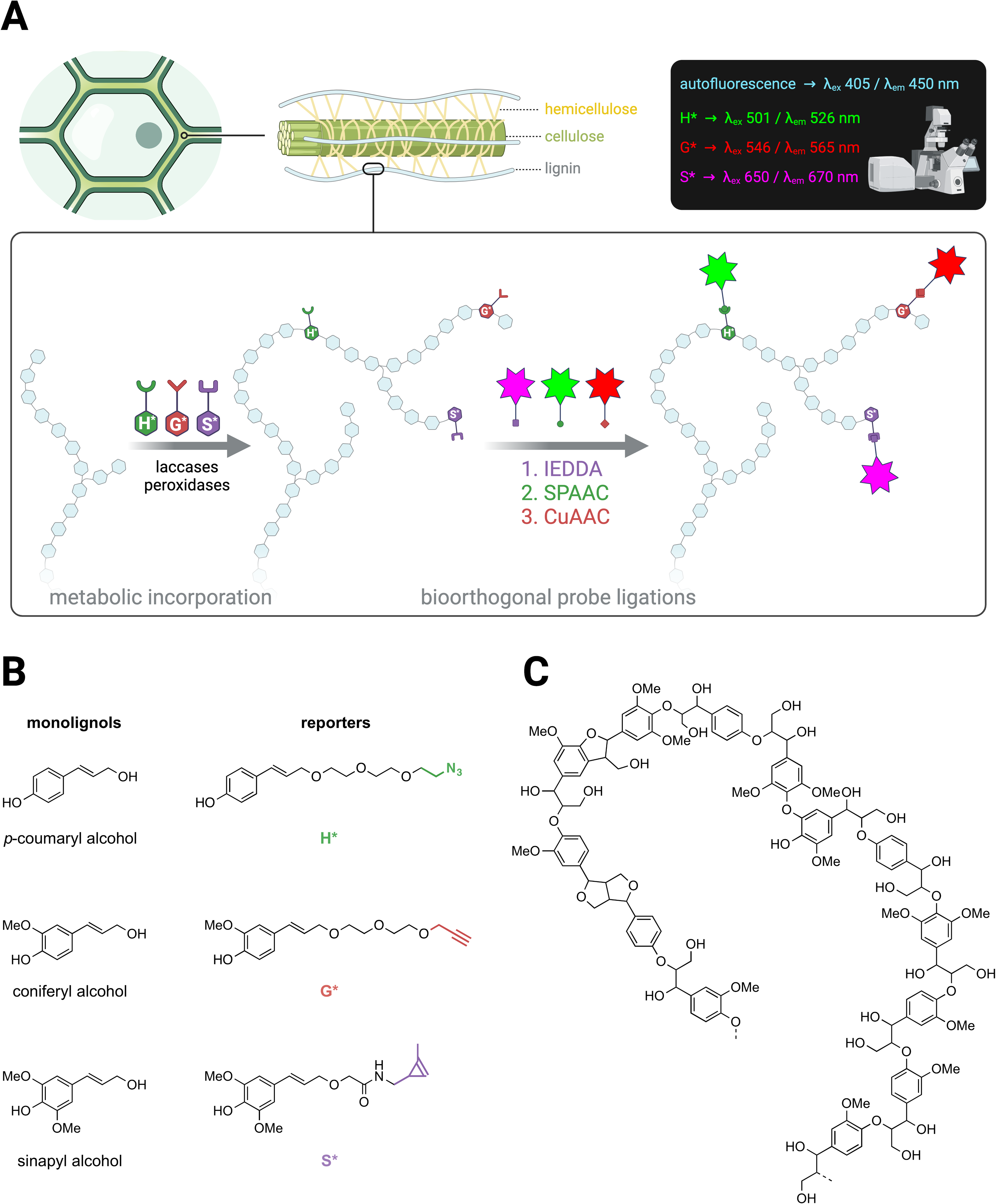
Lignin bioorthogonal triple labeling and chemical reporter structure. **A**) Schematic representation of the lignin biorthogonal triple labelling strategy in plant cell walls. Monolignol chemical reporters H*, G* and S* are oxidized by cell wall located peroxidases and/or laccases and become metabolically incorporated into the growing lignin polymer. Following reporter incorporation, green, red and magenta fluorescent probes are added and become specifically linked to H*, G* and S* reporters, respectively via 3 sequential biorthogonal reactions: Inverse Electron Demand Diels-Adler (IEDDA: S*), Strain Promoted Azide-Alkyne Cycloaddition (SPAAC: H*) and Copper activated Alkyne-Azide Cycloaddition (CuAAC: G*). Spatial localization of fluorophores is analyzed by confocal laser scanning microscopy (CLSM). **B**) Chemical structures of native lignin monolignols (left) and corresponding chemical reporters H*, G* and S* (right). Reporter tags involved in the biorthogonal reaction with the corresponding fluorophores are shown in color: green – azide group, red – alkyne group, magenta – methyl cyclopropene group. **C**) Example of typical lignin oligomer structure.

### 2. Parametric and AI based segmentation

To map the localization of the incorporated monolignol reporters we decided to segment the cell wall into three main zones: the cell corner (CC), the compound middle lamella (CML) consisting of the middle lamella and the primary cell wall, and the secondary cell wall (SCW). Once successfully segmented, pixels can be assigned to the different cell wall sectors and the fluorescence intensity corresponding to each incorporated monolignol reporter determined. Two main strategies can be proposed for the crucial segmentation. Traditional parametric methods are mostly based on the intensity and spatial relationships of pixels, while trainable machine learning methods require the user to “teach” the software which pixel is part of what region. Artificial intelligence (AI) based image analysis are highly complementary to classic parametric segmentation procedures and have strongly emerged as a valuable asset in the microscopy image analysis field over the last few years. For the REPRISAL methodology, we developed a global algorithm that enables the user to implement both parametric and AI segmentation (Figure 1).

For parametric segmentation the initial image was acquired according to nyquist sampling criteria and treated according to the algorithm presented in figure 3. The transmission image is split into 3 fluorescence channels (green, red and magenta) that are transformed into 3 binary masks using the Otsu threshold method (Otsu, 1979). These masks are then merged so that all the labelled lignin is taken into account in the automatic segmentation. A set of transforms is then applied to this new mask including the Distance map (Legland et al., 2016), Voronoï (Legland et al., 2016), Enhance Local Contrast (CLAHE) (Pizer et al., 1987), and Find Maxima transforms (Grishagin, 2015). Binary subtraction operations then generate 3 masks corresponding to the cell corner (CC), the compound middle lamella (CML) and the secondary cell wall (SCW).

**Figure 3.**
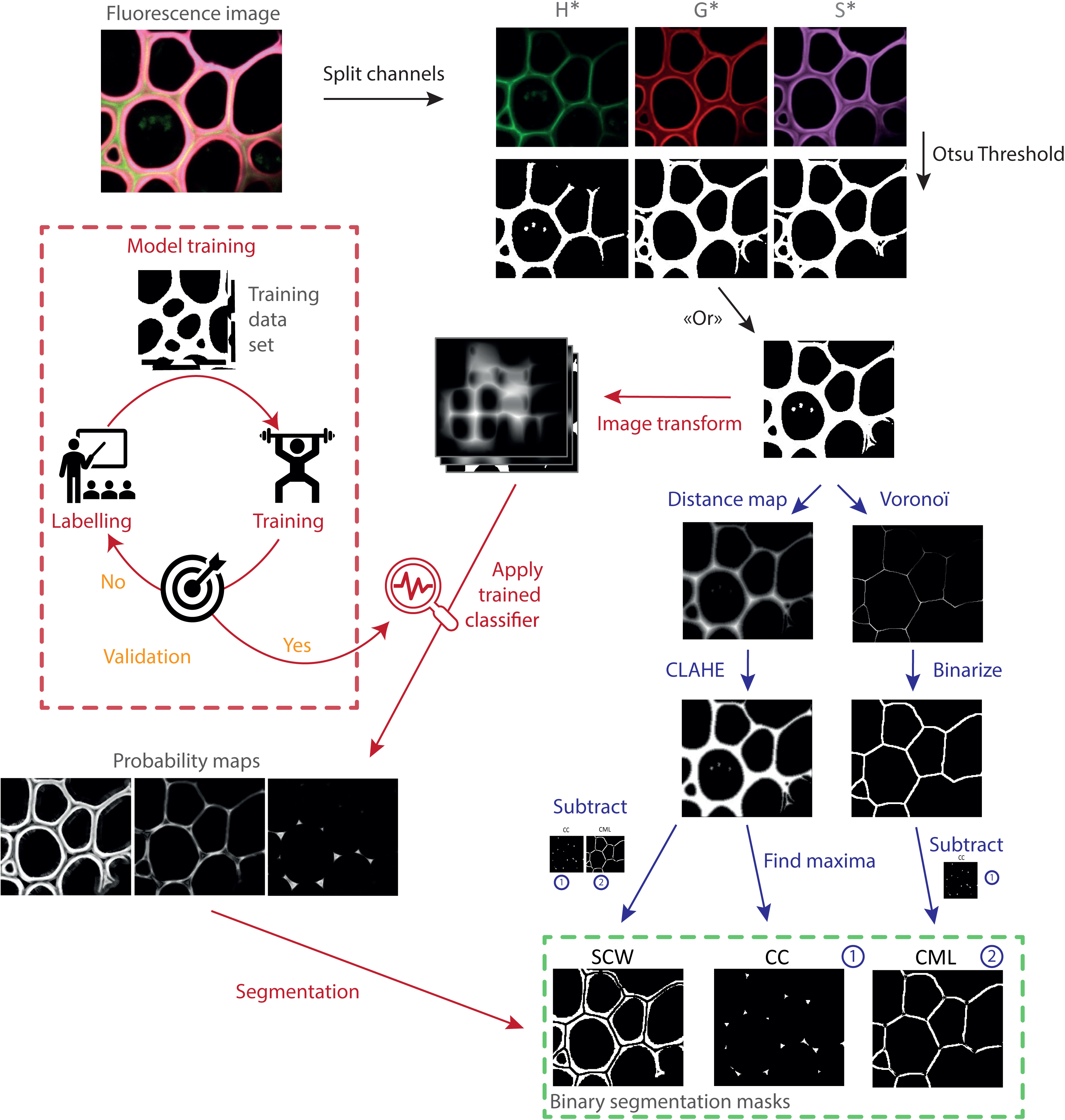
Scheme of the segmentation algorithm. After transforming the images of the individual channels into binary masks and then combining them, the algorithm allows the user to choose between segmentation based on i) morphological parameters (blue path) or ii) learning (red path). Once segmented, a mask is created for the secondary cell wall, the compound middle lamella and the cell corner. It is then applied to each fluorescent channel corresponding to the 3 monolignol reporters (H*, G*, S*). For each fluorescence image, 9 fluorescence intensity maps are thereby created and analyzed.

For AI segmentation and in order to obtain an algorithm easily adaptable to different types of lignin fluorescence images, we chose to keep the first steps of binarization and fusion of binary masks used for parametric segmentation step before implementing the AI segmentation. Images once binarized were manually labeled and then transformed by the WEKA algorithm (“Gaussian Blur”, “Sobel”, “Hessian”, “Differences of gaussians”, “membrane projection” with a membrane patch size of 19, a maximum sigma of 16). Transformed images were then used to train our network (200 initial trees and 4 initialization classes i.e., background, CC, CML and SCW). Once the training was validated on training images, it was applied to a new set of validation images. The classifier we obtained was applied for batch analysis and provides probability maps of each pixel belonging to each category. Probability maps were then converted into binary masks using auto-threshold and despeckle.

To evaluate the efficiency of the developed algorithm, we applied it to confocal microscopy images of Arabidopsis floral stem cross-sections in which lignin was labeled by the bioorthogonal triple marking strategy (figure 4). Visual inspection of the different areas confirms the efficiency of the segmentation. The form and location of the different objects obtained correspond to expectations: cell corners (CC) are triangular in shape and located at the interstices where 3 or more cells are in contact, the compound middle lamella (CML) corresponds to a line formed at the junction between 2 cells, and the secondary cell wall (SCW) occupies the rest of the total cell wall zone. Overall, the parametric and AI segmentation methods give very similar results although some differences can be detected. The parametric method appears to be more accurate for fine structures whereas the AI method recognizes better the cell corners and the middle lamella and follows more closely the curves of the cells. Nevertheless, the percentage distribution of fluorescence intensity values between the three cell wall zones is similar for both methods showing that they can be used indifferently (Table 1). All subsequent results presented in this paper were obtained using parametric segmentation.

**Figure 4.**
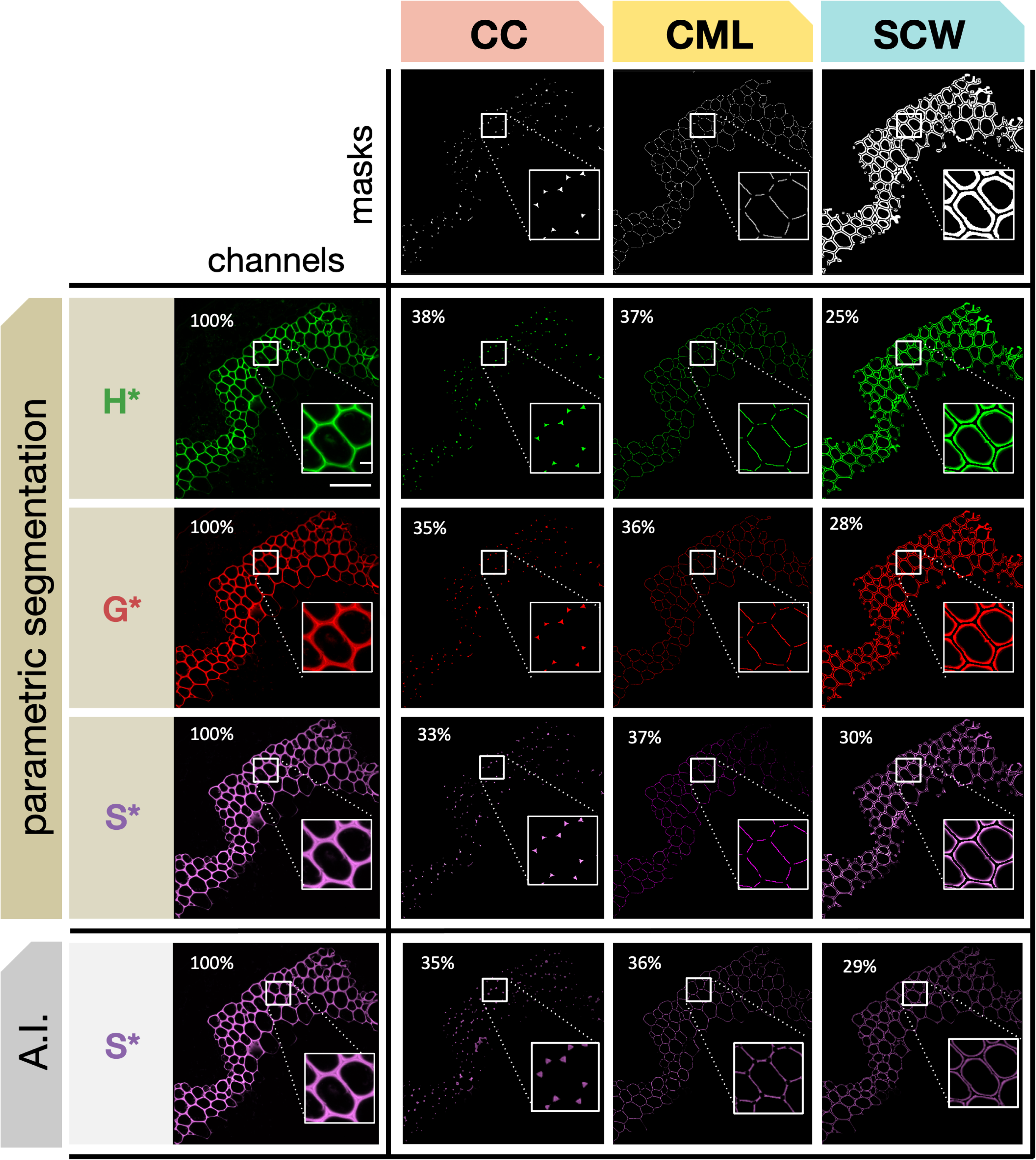
Automatic segmentation of different cell wall zones. The 3 masks (CC, CML, SCW) are applied to cross-sections of Arabidopsis stems in which lignin has been labelled by the triple bioorthogonal strategy. Masks are applied to the three fluorescence channels corresponding to H*, G* and S* reporters. Percentage values indicate the relative proportion of total reporter signal intensity determined for each cell wall zone by parametric segmentation (H*, G* and S*) or artificial intelligence (AI) segmentation where the value for S* is shown for comparison (see Table 1 for corresponding H* and G* values).

**Table 1.** Comparison of fluorescence signal distribution in different cell wall zones determined by parametric segmentation (PS) and artificial intelligence (AI) segmentation. H*, G*, S* = monolignol reporters, CC = cell corner, CML = compound middle lamella, SCW = secondary cell wall. ^1^Raw fluorescence signal intensity.

| Reporter | CW region | Intensity<br>PS<sup>1</sup> | Intensity<br>AI<sup>1</sup> | % Intensity<br>PS | % Intensity<br>AI |
| --- | --- | --- | --- | --- | --- |
| H* | CC | 1807 | 1805 | 38 | 40 |
|  | CML | 1767 | 1631 | 37 | 36 |
|  | SCW | 1172 | 1078 | 25 | 24 |
| G* | CC | 515 | 543 | 35 | 38 |
|  | CML | 530 | 503 | 36 | 35 |
|  | SCW | 416 | 399 | 28 | 28 |
| S* | CC | 1147 | 1214 | 33 | 35 |
|  | CML | 1277 | 1252 | 37 | 36 |
|  | SCW | 1016 | 983 | 30 | 29 |

### 3. Ratiometric analysis

During the bioorthogonal reactions each incorporated monolignol becomes linked to a different fluorophore. Since different fluorophores do not give the same response in terms of the quantity of photons emitted for the same number of incorporated reporters it is not possible to directly compare the amounts of the different reporters incorporated into the three cell wall zones (Schwartz et al., 2002). To get around this problem and obtain quantitative data that can be statistically treated to provide information about differences in reporter incorporation we developed a ratiometric approach. The rationale for this is based on the fact that the ratios between the fluorescence signal of the different reporters do not change for the same biological sample/condition, thereby allowing the comparison of two or more cell wall zones. While this approach does not allow direct comparisons of the absolute quantity of an incorporated reporter, it does make it possible to determine whether there are changes in the relative incorporation of monolignols reporters incorporated into the cell wall zones of different samples.

In the ratiometric approach relative monolignol reporter incorporation is calculated by two methods RM1 & RM2 (ratiometric method 1 & ratiometric method 2) that provide complementary information (Fig. 5). RM1 evaluates the relative ***distribution*** of a given reporter in the different segmented cell wall regions (Fig. 5B). Here, the fluorescence intensity of a given reporter (H*, G*, S*) in a given cell wall region (CC, CML, SCW) is divided by the total fluorescence intensity of that reporter in all three cell wall zones and expressed as a percentage (e.g., (H*CCintensity)/(∑H*). RM2 evaluates the relative ***proportion*** of each reporter in a given cell wall zone compared to the other reporters (Fig. 5C). Here, the fluorescence intensity of a given reporter (H*, G*, S*) in a given cell wall region (CC, CML, SCW) is divided by the total fluorescence intensity of all three reporters in this particular cell wall zone and expressed as a percentage (e.g., (H*CCintensity)/(∑CCintensity). Our results (Fig. 5) illustrate the interest of the combined chemical reporter strategy, segmentation and ratiometric analysis approach for mapping developmentally-related lignification in plants. In this experiment we compared the capacity of different cell wall regions to incorporate monolignol reporters at different developmental stages by analyzing floral stem cross-sections sampled at three different heights from 7-week-old plants. Visual inspection of merged confocal microscopy images in the three cross-sections (Fig. 5A, Supp data 2 reveals that the 3 reporters are incorporated into fiber cell walls at all developmental stages from the “youngest” (23 cm) to the “oldest” (1 cm). Overall, the amount of signal increases during development reflecting reporter incorporation into the secondary cell wall. Further interpretation of these images by visual inspection is difficult due to the huge amount of data that they contain and it is therefore necessary to apply the segmentation approach.

**Figure 5.**
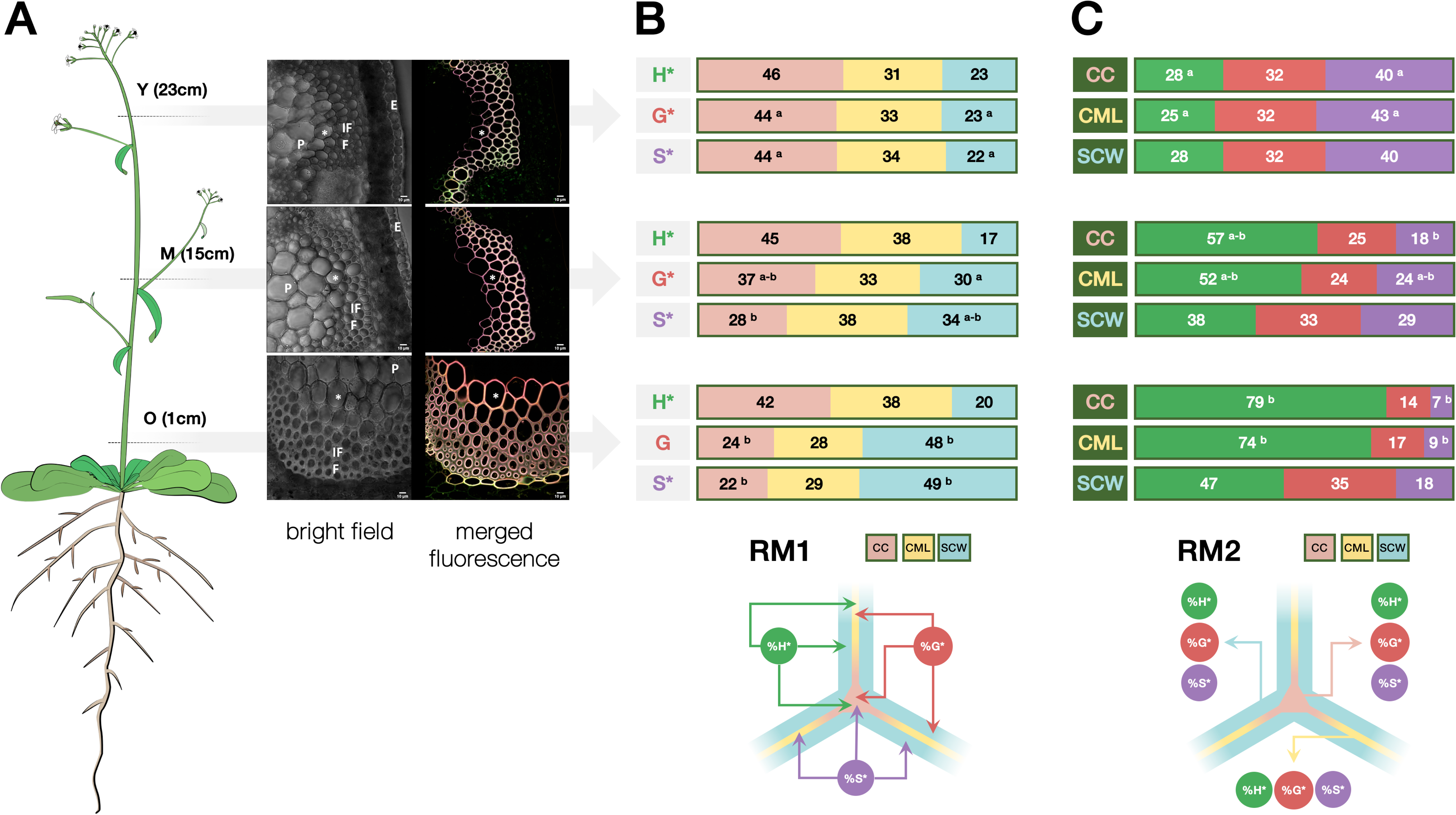
Monolignol reporter incorporation profiles in Arabidopsis stem fiber cell walls. **A**) CLSM images of interfascicular fiber bundles in Arabidopsis stem cross-sections, (i) cartoon of plant indicating regions used to prepare cross-sections, Y = “young” stem region (23 cm from the base of the floral stem), M = “medium” stem region (15 cm from the base), O = “old” stem region (1 cm from the base), total stem height = 30 cm; ii) bright-field image of stem region showing fibers analyzed, IFF = interfascicular fibers, P = pith, E = epidermis, * indicates the same cell shown in the merged image, scale bar = 10 μ; merged green (H*/DBCO-PEG_4_-Rhodamine Green), red (G*/Azide-fluor 545) and magenta (S*/tetrazine-Cy5) channels revealing monolignol reporter incorporation in fiber cell wall lignin during metabolic feeding, * indicates the same cell shown in the bright field image, scale bar = 10 μ. **B**) Relative distribution of each reporter incorporated into different fiber cell wall zones in Y, M and O stem cross sections analyzed by ratiometric method 1 (RM1), CC = cell corner, CML = compound middle lamella, SCW = secondary cell wall; figures represent the percentage of total signal for a given reporter incorporated into the different wall zones, different letters indicate significantly different values (p-value = 0.05) between Y, M and O sections, the absence of a letter indicates that differences are not significant. C) Relative contribution of all reporters to total signal in a given cell wall zone analyzed by ratiometric method 2 (RM2), CC = cell corner, CML = compound middle lamella, SCW = secondary cell wall; figures represent the percentage contribution of each reporter’s signal to total signal intensity in each cell wall zone, different letters indicate significantly different values between Y, M and O sections, the absence of a letter indicates that any differences are not significant.

In young stem sections the use of RM1 (Fig. 5B) indicated that the relative distribution of the total signal in the different cell wall zones is very similar for all 3 reporters with the majority of signal being incorporated into the CC, followed by the CML and SCW (CC: 44 – 46 %; CML: 31 – 34 %; SCW: 22 – 23 %). While this distribution pattern did not evolve over time for the H* reporter clear changes could be observed for G* and S* reporters. The relative proportions of total G* and S* signal significantly decreased in the CC during development accompanied by an increase in the percentage of total signal observed in the SCW. No changes were observed in the relative distribution of signal in the CML. The differences in the relative distribution of the G* and S*, but not H* reporters suggest the existence of developmentally related changes in the capacity of the different cell wall zones to incorporate reporters. The increased capacity of the SCW to incorporate G* and S*, but not H* reporters, could indicate differences in the cell wall machinery that specifically favorize the incorporation of these two reporters compared to H* reporters. If the increased incorporation capacity was simply due to the fact that the secondary cell wall zone is bigger in older sections then one would also expect a similar increase in H* incorporation. The fact that this is not so argues for the existence of a specific mechanism favorizing G* and S* incorporation.

Analysis of the same images using RM2 (Fig. 5C) shows that the contribution of H* to the total signal in the CC increases approximately fourfold (28 % to 79 %) in older stem sections compared to younger stem sections. Since the relative distribution of total H* signal measured in the CC remains relatively stable (42 - 46 %) during development (Fig. 5B), this would suggest that the amounts of G* and/or S* incorporated into the CC decrease during development. This hypothesis is supported by the observations that i) the relative proportions of the total G* and S* signal observed in the CC decrease (Fig. 5B) and ii) that the relative contributions of S* to the total signal also decreases (Fig. 5C). The G* contribution also decreased, but not significantly. The fact that the relative contribution of the S* reporter is more heavily affected (sixfold reduction) than the G* reporter (twofold reduction) suggests that older CCs are less able to incorporate S* reporters.

Analysis of the CML using RM2 gave rise to a similar result, i.e., an approximately threefold increase in the contribution of the H* signal to the total signal in older stem sections (Fig. 5C), associated with a decrease in the relative contributions of G* and S* reporters suggesting that these latter are less easily incorporated in older CMLs compared to H* reporters. Once again, the effect was more marked for S* compared to G*. In comparison to the CC and CML regions, the developmental pattern of reporter incorporation in the SCW showed a more complex pattern. RM2 (Fig. 5C) indicated that the contribution of H* to the total signal in this zone increased while the S* contribution decreased as previously observed for the CC and CML regions. In contrast, the relative G* contribution remained unchanged. Although the S* contribution to total signal in the SCW decreased in older stem sections as for the CC and CML zones the extent of the reduction was far less (twofold) compared to a six-fold reduction (CC) and four-fold reduction (CML). These results are coherent with those obtained by RM1 (Fig. 5B) showing that the relative proportion of the G* and S* total signals increased in the SCW.

Overall, these results indicate that while H* reporters are the most readily incorporated form into all three cell wall regions at all of the developmental stages examined, there is an increasing tendency to incorporate both G*, and especially S* reporters into the SCW of older stem sections. It is possible that these changes are functionally related to the developmentally-related occurrence and distribution of lignin-related peroxidases/laccases recently reported by the group of Lacey Samuels (Hoffmann et al., 2020).

In conclusion, these results clearly demonstrate the potential of the REPRISAL methodology that combines chemical reporter + segmentation + ratiometric analysis approaches for mapping developmental lignification in Arabidopsis.

### Lignification mapping in the Arabidopsis *prx64* mutant

Having demonstrated the interest of REPRISAL in WT plants we decided to use this approach to investigate whether it would be capable of revealing differences in the lignification profile of a lignin mutant compared to WT. For this we chose the Arabidopsis peroxidase *prx64* mutant (AT5g42180). Several studies show that the AtPRX64 protein is associated with lignification; it is involved in Casparian strip lignification (Lee et al., 2013) and the AtPRX64-mCherry fusion protein becomes localized in the lignified cell corners and middle lamella of Arabidopsis fiber cells in a developmentally-related pattern (Yi Chou et al., 2018; Hoffmann et al., 2020). The AtPRX64 promoter is also active in both xylary and interfascicular fibers (Smith et al., 2017). Despite this apparently clear link between PRX64 and lignification, the stem lignin content has not been investigated in this mutant and we therefore decided that it would be a good target to evaluate the efficiency of our methodology. A visual inspection of both UV autofluorescence and merged triple labeling images from WT and mutant plants suggested that lignification was affected in the mutant (Supp Data 3).

While comparison of lignin UV autofluorescence images showed no differences between WT and mutant young (Y) sections, a clear difference was observed in the color of sections obtained by triple labeling. Merged images showed that the great majority of WT fiber cell walls were colored yellow with some cell walls closer to the pith showing a pinker color. In contrast, mutant cell walls were uniformly more red-pink in color indicating changes in the relative incorporation of different reporters. This observation was confirmed by ratiometric analysis (see below) and suggests that the absence of a functional PRX64 protein modifies the monolignol oxidizing capacity of the cell wall at this developmental stage resulting in a preferential uptake of G* reporter to H* reporter. Further examination of the merged image also showed little labelling in the SCW region suggesting that the development of this cell wall layer is negatively affected in the mutant compared to WT. Once again, this observation was confirmed by the ratiometric analysis (below). Differences could also be observed in the color of M sections obtained by triple labeling; the more orange (less pink) color of mutant cell walls is coherent with ratiometric analysis indicating increased H* and reduced S* incorporation compared to WT. In O sections, clear differences between WT and mutant fiber cell walls were observed for both UV and triple labelling images. The first technique showed that lignin was uniformly distributed throughout all of the WT cell wall layers including the thick S2 secondary cell wall characteristic of fibers at this developmental stage. In contrast, UV autofluorescence revealed that lignin appeared to be limited to the S1 layer of the secondary cell wall and was not detectable (by this technique) in the S2 layer. Triple labelling also revealed important differences in the lignification process between the WT and *prx64*. In partial agreement with the UV images, triple labelling suggested that secondary cell wall development was negatively affected in the mutant as shown by the generally thinner secondary cell walls. In addition, the color differences also indicated changes in the relative incorporation of different monolignol reporters between WT and mutants with preferential incorporation of H* reporters and decreased S* incorporation in the mutant as confirmed by ratiometrics.

The figure 6 shows the comparison of the application of RM1 for the G* reporter (Fig. 6A) and RM2 for CC (Fig. 6B) in the *prx64* mutant compared to WT as an example of the information that can be obtained by our approach. Corresponding data for H* and S*, and CML and SCW are given in Supp Data 4. Analysis by RM1 indicates significant differences in the relative distribution of total signal for all 3 reporters between mutant and WT samples for the Y (young) stage, but not M and O stages. In Y samples, significantly higher amounts of all reporters are incorporated in the mutant CML compared to WT. At the same time, significantly lower amounts of H* and G* (but not S*) are incorporated into the SCW. No significant differences are observed for the CC zones. These results would suggest that the capacity of the CML and SCW, but not CC, to incorporate monolignol reporters is modified in the *prx64* mutant compared to WT only in Y samples. This is intriguing on several levels; firstly, studies by the group of Lacey Samuels (Yi Chou et al., 2018; Hoffmann et al., 2020) have shown that the PRX64 protein is localized to the CC region and it could therefore be expected that the mutation would preferentially induce a modification in this zone rather than the others. Secondly, the same group showed that the protein was not detected in stage 1 (young) fiber CC, but only in the developmentally more advanced stage 2 and stage 3 CC. However, our plants were grown in short day conditions compared to those of Samuels and co-workers, similarly, our Y, M and O samples were obtained from 30-cm high floral stems of 8-week-old plants (c.f. 18 cm high, 5-week-old plants in the other study). As a result, our Y, M and O samples are not directly comparable to the stage 1-3 samples used by Samuels and it is possible that differences between the developmental stages analyzed could partly explain the observed discrepancy. Another interesting observation concerns the relative *increase* of all 3 reporters incorporated in the mutant CML zone of the mutant compared to WT since it would be expected that the KO of a lignin peroxidase gene would lead to a reduced reporter incorporation. However, it is also possible that the mutation provokes modifications in the expression profiles of other *PRX* and/or *LAC* genes thereby changing the overall incorporation capacity of the different cell wall regions. Future transcriptomics could help to clarify this point.

**Figure 6.**
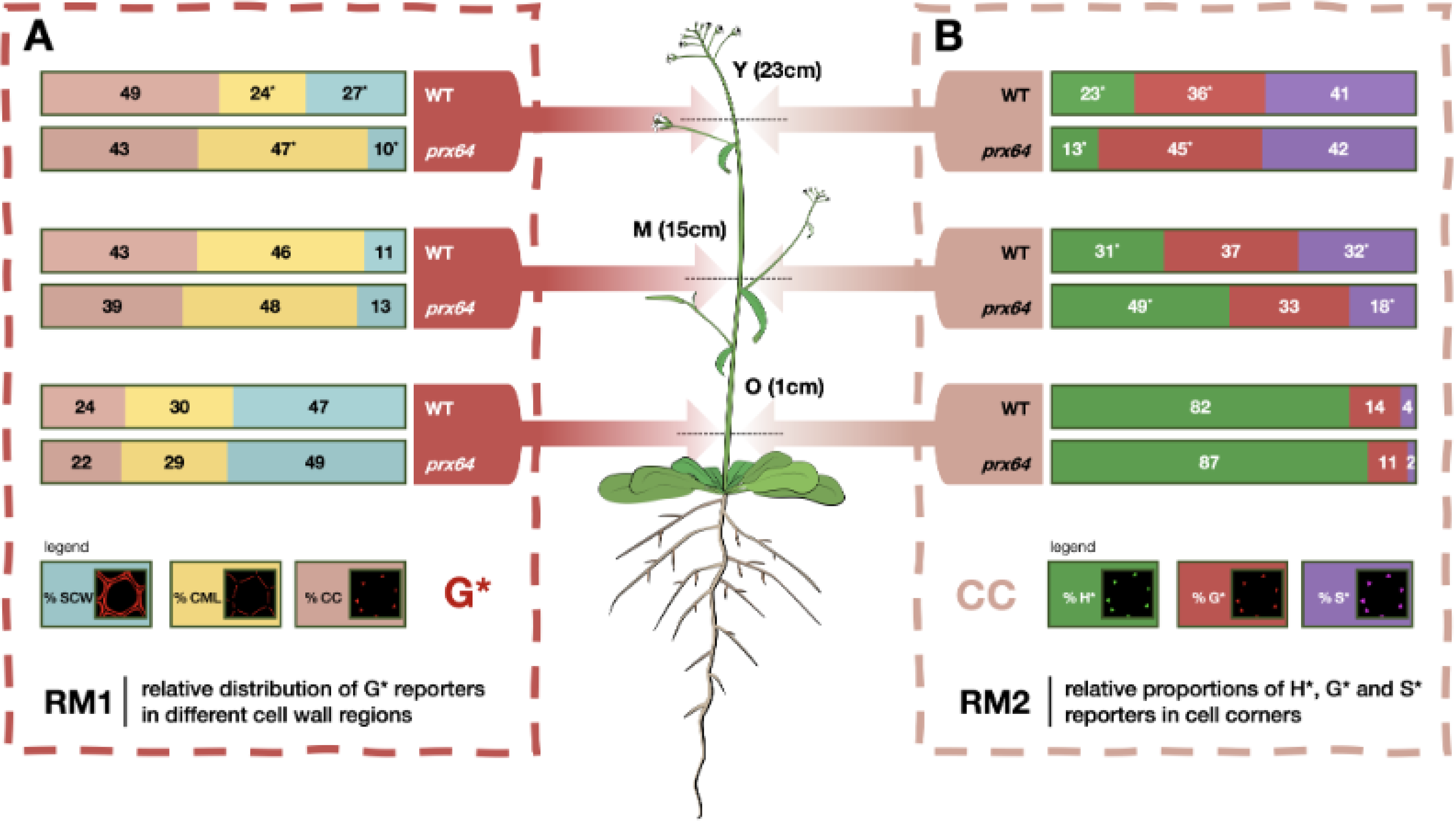
Example of a comparison of monolignol reporter incorporation profiles in WT and *prx64* mutant Arabidopsis stem fiber cell walls. **A**) Relative distribution of G* reporter incorporated into different fiber cell wall zones in Y, M and O stem cross sections analyzed by ratiometric method 1 (RM1), CC = cell corner, CML = compound middle lamella, SCW = secondary cell wall; figures represent the percentage of total G* signal incorporated into the different wall zones, different letters indicate significantly different values between WT and mutant plants for a given stem height (Y, M, O), the absence of a letter indicates that differences are not significant. **B**) Relative contribution of all reporters to total signal in the cell corner (CC) analyzed by ratiometric method 2 (RM2); figures represent the percentage contribution of each reporter’s signal to the total CC signal intensity, different letters indicate significantly different values between WT and mutant plants for a given stem height (Y, M, O), the absence of a letter indicates that differences are not significant. Corresponding figures for H* and S* probes (RM1) and CML and SCW zones (RM2) are shown in **Supplementary Data 4**.

Analysis by RM2 (Fig. 6, Supp Data 4) shows that the relative proportions of the 3 reporters in each cell wall region also change in the mutant compared to WT. A significantly lower and higher proportion of H* and G* reporters, respectively, are incorporated in all 3 cell wall regions of the mutant compared to WT in Y samples; the relative proportions of S reporters remain unchanged. In M samples, the trend is inversed with a significantly higher proportion of H* reporter being incorporated in the CC and CML regions of the mutant compared to WT. In the mutant SCW, although the relative proportion of H* increases, the difference is not significant. The increase of H* in the SCW layer is not significant. The analysis also shows that the relative proportions of S* reporter incorporated into all 3 regions increase in the mutant compared to WT; G* reporter proportions are not significantly affected. Finally, in O samples, the only significant change concerns the decrease in S* reporter incorporation in the mutant SCW region compared to WT.

Taken together, our results show that both the RM1 and RM2 analyses, when combined with the bioorthogonal click chemistry and segmentation methodologies reveal significant complex changes to the spatial lignification profile in the *prx64* mutant compared to WT. Such an observation firstly illustrates the complexity of lignification at the cell wall layer scale. A complete understanding of this process is complicated because of the number of different actors that are involved. In addition to the different redox enzymes such as laccases and peroxidases it is also necessary to take into account other factors such as the supply of H_2_O_2_ for peroxidases as recently demonstrated (Hoffmann et al., 2020). Secondly, our results also illustrate the enormous potential of the combined labelling + segmentation + ratiometrics to finely dissect changes in the lignification capacity of cell wall mutants, including those where classical approaches have previously failed to detect differences.

### Lignification mapping in different plant species

To ensure the robustness of REPRISAL, and to evaluate its interest for mapping lignification we applied it to three other plant biomass species: poplar, flax and maize (Figure 7, Table 2). Our results firstly confirm that the bioorthogonal triple-labelling strategy is applicable to different plant species as previously observed (Simon et al., 2018). Secondly, they show that the developed macro is capable of successfully segmenting cell walls from different species into the 3 different zones (CC, CML and SCW) previously analyzed in Arabidopsis. Closer examination of the % distribution of each reporter in the different cell wall zones revealed a number of potentially interesting differences in lignification in the different species.

**Figure 7.**
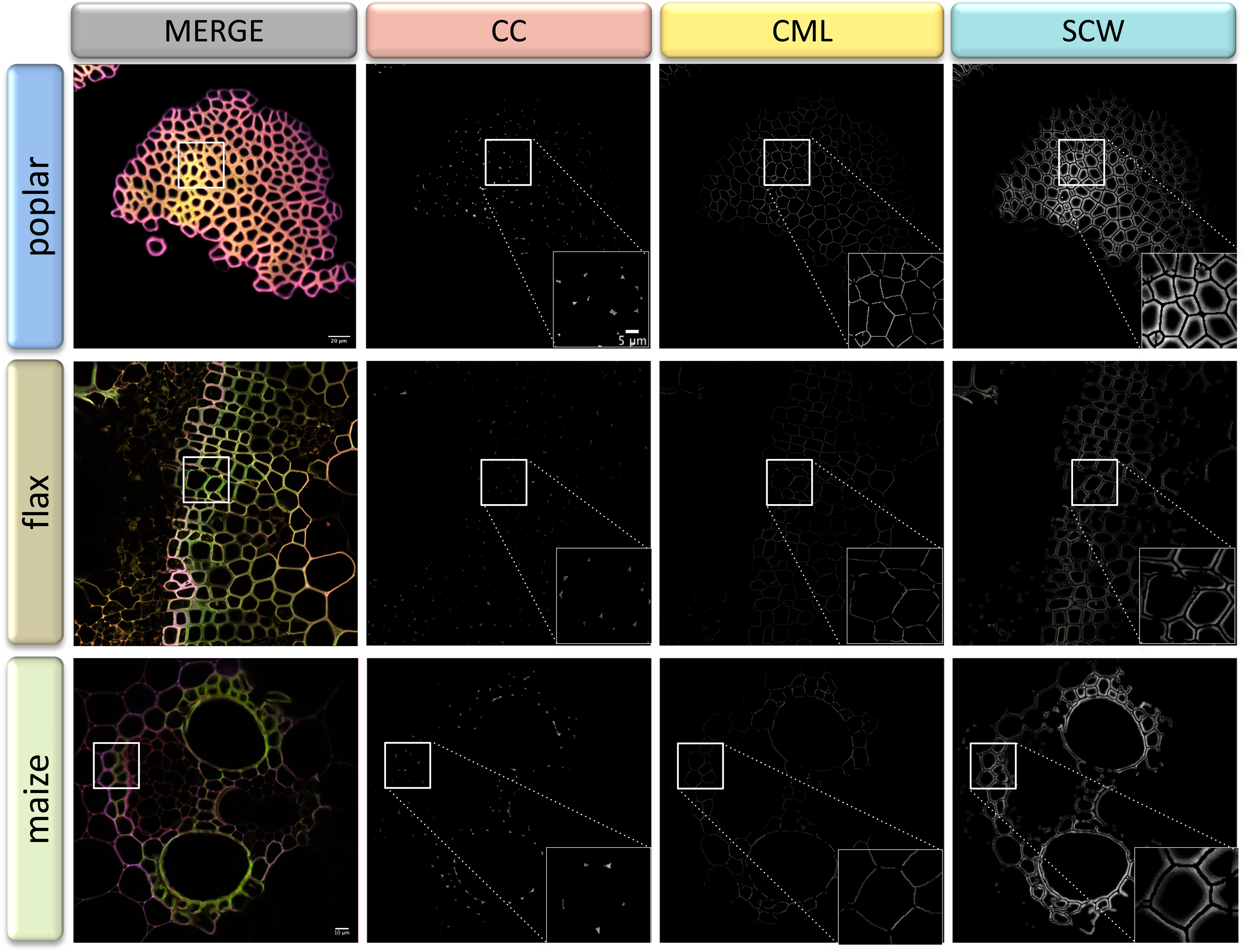
Automatic segmentation applied to different plant species. Top line: poplar stem bark fibers, middle line: flax stem xylem, bottom line: maize stem vascular bundle. First column: merged CLSM image of green, red and magenta fluorescence channels, second – fourth columns: cell wall zone segmentation for CC (cell corner), CML (compound middle lamella) and SCW (secondary cell wall). All samples labelled by the triple lignin biorthogonal chemical reporter strategy with H*, G* and S* monolignol reporters. Square frames indicate the sample region magnified (bottom RHC, columns two – four).

**Table 2.** Comparison of % total reporter distribution in different cell wall zones for different plant species determined by RM1. H*, G*, S* = monolignol reporters; CC = cell corner, CML = compound middle lamella, SCW = secondary cell wall; red values = highest %, green values = lowest %.

| Reporter | CW region | % Intensity<br>Arabidopsis | % Intensity<br>poplar | % Intensity<br>flax | % Intensity<br>maize |
| --- | --- | --- | --- | --- | --- |
| H* | CC | 38 | 31 | 31 | 35 |
|  | CML | 37 | 28 | 37 | 35 |
|  | SCW | 25 | 40 | 32 | 30 |
| G* | CC | 35 | 48 | 35 | 36 |
|  | CML | 36 | 23 | 35 | 35 |
|  | SCW | 28 | 29 | 29 | 29 |
| S* | CC | 33 | 42 | 29 | 34 |
|  | CML | 37 | 25 | 39 | 37 |
|  | SCW | 30 | 32 | 32 | 29 |

In Arabidopsis, poplar and maize, data was obtained on cell walls in functionally similar “support tissues” consisting of interfascicular fibers (Arabidopsis), bark sclerenchyma fibers (poplar) and vascular bundle sclerenchyma (maize). Despite their biologically similar roles, the cell walls of poplar fibers incorporated the majority of the H* reporter into the SCW in comparison to Arabidopsis and maize where the reporter was preferentially incorporated into CC lignin. A similar difference was observed for S* reporter that was preferentially incorporated in the CC for poplar compared to the CML for the other 2 species. In contrast, the distribution of incorporated G* reporter was similar in all 3 species. While such differences can most probably be related to the developmental stage of the cells/tissues analyzed the results clearly demonstrate the potential of our approach for large-scale and detailed mapping of cell wall lignification at the cell wall layer level in different species. Our results on flax also show that the approach can be successfully applied to lignifying cell walls in other tissues (xylem).

## Conclusions

In this paper we have combined three different methods (chemical reporter + segmentation + ratiometric analysis) to generate a robust methodology for analyzing lignification in plants that we have termed REPRISAL for **REP**orter **R**atiometrics **I**ntegrating **S**egmentation for **A**nalyzing **L**ignification. Central to this approach is the development of a new segmentation method allowing automatic quantification of the fluorescence signal resulting from lignin bioorthogonal triple labeling in different cell wall zones. Pixel segmentation, i.e. their distribution between cell corners, compound middle lamella and secondary cell wall is determined via 2 methods, one parametric and the other based on artificial intelligence. As we have shown in Figure 4, these two methods produce very similar results, and while leading to the same biological conclusions, are not identical. On structures of such complexity, the labeling of pixels by 2 different human experts would also be expected to give slight variations and, in this case, it would be preferable that the analysis of the data set be done by a single expert for the sake of reproducibility. The two experts can also individually analyze all the data before subsequently comparing their interpretations. The same rigor must also be applied when using automated systems such as those proposed and it is therefore important not to mix segmentation methods during analysis even if any systematic bias is most likely minimal.

In the majority of this paper, we have focused on the use of parametric segmentation to demonstrate its applicability to different conditions and plant models. We chose to validate this method because it is the one which is the least “flexible” in terms of use. While effective parametric segmentation is usually simple and quick to use when applied to a given type of image, the determined parameters depend upon the size of the pixels, and on the size and shape of the structures. It is therefore often best suited for use at a given magnification, or even to a species or a plant tissue and requires a high expertise in order to be able to apply it to new types of samples. Nevertheless, as we have demonstrated our parametric segmentation remains adaptable to a large number of situations and is, if possible, the method that should be privileged, because of a greater execution speed (seconds compared to minutes per image analyzed). On the other hand, the WEKA based (AI) method is in essence adaptable to any sample by training it on new sets of samples representative of the condition to be analyzed. Furthermore, the number of transforms performed by the presented algorithm was deliberately limited to reduce computation time. Adding additional complementary filters such as, “kuwahara”, “Gabor” or “neighbors” could further improve the ability of the algorithm to distinguish different cell wall areas. Nevertheless, the inclusion of a batch mode in our plugin makes it possible to launch the analysis on a large number of images overnight in order to optimize the analysis pipeline.

Our algorithm combines the advantages of these two methods by first proposing to test a parametric method on the samples. If this is validated, it can then be applied automatically on a large number of images. Otherwise, the user is offered to test a WEKA reference training set which can in turn be automated or used to initiate a new complementary training phase. Our overall segmentation methodology therefore benefits from the speed of execution of a parametric segmentation when possible and from the flexibility of WEKA for samples that are too different from our training sets.

The second essential element that contributes to the potential of REPRISAL was the development of a ratiometric approach. Such an approach is necessary because different fluorophores emit different amounts of photons and the measured fluorescence intensities obtained with different fluorophores do not therefore accurately reflect the amount of reporter incorporated. Our approach is based upon two robust and easily interpretable intensity ratios; RM1 evaluates the relative ***distribution*** of a given reporter in the different segmented cell wall regions while RM2 evaluates the relative ***proportion*** of each reporter in a given cell wall zone. These methods made it possible to extract quantitative information from the complex images generated by the triple monolignol reporter labeling strategy that could then be statistically analyzed to generate useful biological information as discussed above.

The final element necessary to REPRISAL was the implementation of the previously developed triple labelling strategy (Simon et al., 2018). When applied to the Arabidopsis *prx64* mutant, REPRISAL was able to detect statistically significant differences in the lignification pattern compared to WT even though a simple visual evaluation was unable to this. In addition to confirming that the AtPRX64 protein does indeed play a role in cell wall lignification in stems, this observation also confirms previous results from our group showing that the chemical reporter approach is able to reveal mutant phenotypes undetectable by classical cell wall analyses (Baldacci-Cresp et al., 2020a).

In this paper we focused on mapping lignification and we therefore used 3 monolignol chemical reporters. However, as previously demonstrated (Simon et al., 2018), the triple labeling strategy is also compatible with monosaccharide reporters and it should therefore be possible to exploit REPRISAL for mapping the distribution of other cell wall polymers including pectin and hemicelluloses. A number of monosaccharide reporters have been successfully used in mono-labelling approaches by other groups and could be used (Anderson et al., 2012; Dumont et al., 2016; Zhu et al., 2016).

Finally, it would also be interesting to adapt REPRISAL, or at least the segmentation component, to the analysis of cell wall images generated by other imaging techniques in order to multiply the information that can be obtained. Possible targets could include, for example, different histochemical reagents such as basic Fuschin, Safranin, or Auramine O that have been shown to be compatible with fluorescence microscopy (Ursache et al., 2018; Baldacci-Cresp et al., 2020b), as well as antibodies capable of recognizing a wide range of cell wall epitopes (Donaldson and Knox, 2012) and other fluorescence-based imaging techniques (DeVree et al., 2021). Another major direction could be to explore the possibilities of combining REPRISAL-type segmentation with the segmentation resulting from multivariate analyses of vibrational spectroscopic data such as Raman (Gierlinger et al., 2012) or Atomic Force Microscopy (Felhofer et al., 2020). Such approaches could be expected to contribute to our understanding, not only of lignification, but also of cell wall biology in general, thereby paving the way for exciting new discoveries in both fundamental and applied areas.

## Materials and Methods

### Plant material

*Arabidopsis thaliana* (Columbia, Col-0) plants were used for experiments. Seeds were stratified at 4°C in 0.1% Phytagel solution (w/v) for 3 days before being sowning. Plants were grown in growth chambers (GroBank, BB-XXL.3+) under 12-h light cycles (120 µmol photon m^−2^ s^−1^) at 23°C during the day and 20°C during the night. The *atprx64* insertion mutant (SALK_203548C) was purchased from the Nottingham Arabidopsis Stock Centre (NASC)

*Linum usitatissimum* (flax) and *Zea mays* (maize) plants were grown in growth chambers (Angelantoni Life Sciences) at 22°C with a photoperiod of 16hr/8hr day/night. *P. tremula* × *Populus alba* (poplar) plants were grown in a phytotron with a 16-h light/8-h dark photoperiod, 24°C/21°C, with 35–65% hygrometry under white light (HPI Master Plus Philips, metal halide) to maintain a light intensity of 120 µmol m−2 sec−1.

### Sample preparation and reporter labeling

Sample preparation was adapted from the protocol described in Simon et al (2018). Sections of 80µm thickness were made with a vibroslicer (VT-1000S, Leica, Wetzlar, Germany) at different heights (1 cm, 15 cm and 23 cm) from the base of the stem of embedded (3.5 % agarose) samples of flowering stems of *Arabidopsis thaliana* plants (7 weeks). The sections were placed in sterile Murashige and Skoog half-strength (½ MS) solution and stored at 4° C prior to incubation with reporters. Maize, flax and poplar sections were made from the stems of 8-week-old plants.

Bioorthogonal triple labeling of monolignol chemical reporters H*, G* and S* (Fig. 2) was performed as previously described (Simon et al., 2018) and outlined in figure 2. Stem cross-sections were incubated in 300 μl ½ MS containing 5 µM of H*, G* and S* for 20 h in the light at 20 °C (Grobank). Control samples were incubated in 300 μl ½ MS containing 5 µM corresponding untagged natural monolignols. After incubation, samples were extensively washed (4 x ½ MS) prior to labelling with fluorophores via three sequential bioorthogonal reactions performed in the following order: 1) Inverse Electron Demand Diels-Adler (IEDDA: S*), 2) Strain Promoted Azide-Alkyne Cycloaddition (SPAAC: H*) and 3) Copper activated Azide-Alkyne Cycloaddition (CuAAC: G*). Tetrazine Cy5 (Jena Bioscience, Jena, Germany) and Azide Fluor 545 (Sigma-Aldrich, Saint-Louis, Missouri, USA) fluorophores were used at 5 µM; DBCO-PEG4-Rhodamine Green fluorophore (Jena Bioscience, Jena, Germany) was used at 2.5 µM. All reactions were carried out in 300 μl ½ MS for 1 h in the dark and samples washed (4 x ½ MS) between reactions. Following the final reaction, samples were extensively washed (2 x 1⁄2 MS, 5 min, 1 x MeOH 70 %, 60 min, 1 x 1⁄2 MS, 5 min and 3 x 1⁄2 MS, 10 min) to fully remove any remaining free fluorophore.

### Confocal microscopy

Image acquisitions were performed as previously described (Simon et al., 2018). Images from 3 independent biological replicates for each genotype and tissue were acquired. A Nikon A1R confocal equipped with a 60x/1.4 aperture oil immersion objective (Plan APO VC) and the NIS Element AR3.0 software was used (Nikon, Tokyo, Japan). Acquisitions were performed on 4 channels corresponding to i) lignin autofluorescence (ƛex: 405, ƛem: 450/50); ii) H units (ƛex: 488, ƛem: 525/50); iii) G units (ƛex: 561, ƛem: 595/50); iv) S units (ƛex: 561, ƛem: 700/75).

### Segmentation

The graphical user interface has been created as a plugin for Fiji using Jython, a Java implementation of Python. Using this language made it possible to use Fiji methods in the Jython script, thereby enabling the creation of the GUI as well as a smooth interaction with Fiji. Fiji macros were used for both AI and parametric segmentation, allowing the program to run with either type of segmentation. The overall analysis procedure was developed with ImageJ 1.53 and JAVA 1.8.

### Parametric segmentation

Parametric segmentation was performed using a homemade imageJ macro. Briefly, the macro first creates a binary mask for i) secondary cell walls, ii) cell wall corners and iii) compound middle lamellae. The binary mask of each region was applied to each fluorescence channel and fluorescence mean values were extracted for the 9 newly-created images. A recapitulative montage image was then created to quickly estimate segmentation quality. The imageJ macro and sample images are available in the Zenodo repository, http://doi.org/10.5281/zenodo.4809980.

### AI Segmentation

The Machine learning approach is based on the “Waikato Environment for Knowledge Analysis” (WEKA) implemented in ImageJ (Witten et al., 2016). We first defined a classification based on four categories: i) secondary cell wall, ii) cell corners, iii) compound middle lamella and iv) background. Our aim was to provide methods compatible with traditional computers and thus limited the training parameters to gaussian blur, Sobel filter, Hessian, Difference of gaussians and membrane projection with a maximum sigma of 16. These training features allowed unambiguous discrimination between the four classes after appropriate training of a random forest classifier with 200 initial trees. Manual labeling was performed on a pixel-by-pixel basis between the four classes. To avoid overlearning, training was performed on confocal images taken from various species, regions and using both wild type and mutants. Each image is then segmented based on obtained probability maps (80% probability is used as a threshold) and compared with results achieved through manual segmentation. Once the training is achieved on reference images, the classifier is applied to new sets of images to validate the training step.

For more details, please see the result section. The imageJ plugin and sample images are available in the Zenodo repository, http://doi.org/10.5281/zenodo.4809980

### Ratiometric Analysis

Two complementary ratiometric methods (RM) were used: RM1 evaluates the relative ***distribution*** of a given reporter in the different segmented cell wall regions. Fluorescence intensity of a given reporter (H*, G*, S*) in a given cell wall region (CC, CML, SCW) is divided by the total fluorescence intensity of that reporter in all three cell wall zones and expressed as a percentage (e.g., (H*CC intensity)/(∑H*). RM2 evaluates the relative ***proportion*** of each reporter in a given cell wall zone compared to the other reporters. Fluorescence intensity of a given reporter (H*, G*, S*) in a given cell wall region (CC, CML, SCW) is divided by the total fluorescence intensity of all three reporters in this particular cell wall zone and expressed as a percentage (e.g., (H*CCintensity)/(∑CC intensity). Statistical differences (p-value of at least 0.05) between samples were determined by the Student’s t-test and analysis of variance (ANOVA).

## Supplementary data

**Supplementary Data 1:** dataset provided in http://doi.org/10.5281/zenodo.4809980

This dataset contains:

- the algorithm with graphical user interface for imageJ and its installation procedure (“Cell_Wall_Segmentation” and “Cell_Wall_SegmentationTutorial”)

- a folder comprising a classifier and a dataset compatible with the machine learning part of the algorithm: “data and classifier for weka”

- representative images adapted for testing: “representative images”

- the macro corresponding to the parametric segmentation procedure (see imageJ documentation for installation instructions): “parametric_segmentation”

**Supplementary Data 2:** Bioorthogonal lignin triple (H*, G*, S*) labelling in Arabidopsis floral stems.

**Supplementary Data 3:** Bioorthogonal lignin triple (H*, G*, S*) labelling in Arabidopsis WT and *prx64* mutant floral stems.

**Supplementary Data 4:** RM1 and RM2 ratiometric comparison of relative monolignol reporter incorporation in Arabidopsis WT and *prx64* mutant floral stems.

